# Bile acids gate dopamine transporter mediated currents

**DOI:** 10.1101/2021.09.13.459497

**Authors:** Tiziana Romanazzi, Daniele Zanella, Mary Hongying Cheng, Behrgen Smith, Angela M. Carter, Aurelio Galli, Ivet Bahar, Elena Bossi

## Abstract

Bile acids (BAs) are molecules derived from cholesterol that are involved in dietary fat absorption. New evidence supports an additional role for BAs as regulators of brain function. Interestingly, sterols such as cholesterol interact with monoamine transporters (MAT), including the dopamine (DA) transporter (DAT) which plays a key role in DA neurotransmission and reward circuitries in the brain. The present study explores interactions of the BA, obeticholic acid (OCA), with DAT and mechanistically defines the regulation of DAT activity *via* both electrophysiology and molecular modeling. We express murine DAT (mDAT) in *Xenopus laevis* oocytes and confirm that DA induces an inward current that reaches a steady-state at a negative membrane voltage. Next, we show that OCA triggers an inward current through DAT that is Na^+^ dependent and not regulated by intracellular calcium. OCA also inhibits the DAT-mediated Li^+^ leak current, a feature that parallels DA action and indicates direct binding to the transporter. Interestingly, OCA does not alter DA affinity nor the ability of DA to promote a DAT-mediated inward current, suggesting that the interaction of OCA with the transporter is non-competitive, in regard to DA. The current induced by OCA is transient in nature, returning to baseline in the continued presence of the BA. To understand the molecular mechanism of how OCA affects DAT electrical activity, we performed docking simulations. These simulations revealed two potential binding sites that provide important insights into the potential functional relevance of the OCA-DAT interaction. First, in the absence of DA, OCA binds DAT through interactions with D421, a residue normally involved in coordinating the binding of the Na^+^ ion to the Na2 binding site (Borre et al., 2014;Cheng and Bahar, 2015). Furthermore, we uncover a separate binding site for OCA on DAT, of equal potential functional impact, that is facilitated through the residues DAT R445 and D436. This binding may stabilize the inward-facing open (IF*o*) state by preventing the re-formation of the IF gating salt bridges, R60-D436 and R445-E428, that are required for DA transport. This study suggests that BAs may represent novel pharmacological tools to regulate DAT function, and possibly, associated behaviors.

## 1 Introduction

Bile acids (BAs) are amphipathic molecules derived from cholesterol that are primarily synthesized in the liver and stored in the gallbladder. Upon food consumption and transit, BAs are released into the duodenum (Mertens et al., 2017) where their main physiological role is solubilization and absorption of dietary fat. Administration of BAs has been developed into successful therapies for the treatment of liver and gallbladder pathologies, such as non-alcoholic steatohepatitis and cholelithiasis (Cruz-Ramon et al., 2017;Li and Chiang, 2020). When administered orally, they exhibit favorable bioavailability. They are readily absorbed through the portal vasculature and distributed throughout the body (Kiriyama and Nochi, 2019).

Receptors for BAs are present throughout the brain (Maruyama et al., 2002;Kawamata et al., 2003;Keitel et al., 2010;Huang et al., 2015;Perino et al., 2021) and evidence exists for their synthesis directly in the CNS (McMillin and DeMorrow, 2016;Mertens et al., 2017;Kiriyama and Nochi, 2019). This raises the possibility of physiologic roles for BAs other than acting as an adjuvant in fat absorption, such as regulators of CNS activity. Consistent with this idea, BAs have been implicated in the modulation of CNS proteins such as NMDA, GABA_A_ and M_3_ muscarinic receptors (Raufman et al., 2002;Schubring et al., 2012), the activation of neuronal ion channels (Wiemuth et al., 2014;Kiriyama and Nochi, 2019) and stimulation of the release of neuroactive peptides such as GLP-1 (Flynn et al., 2019;Chaudhari et al., 2021;Fiorucci et al., 2021). BAs pass the Blood-Brain Barrier (BBB) through passive diffusion, as well as through active membrane transporters, and BA levels in the brain have been correlated to circulating levels (Reddy et al., 2018;Kiriyama and Nochi, 2019). As a result, the number of studies proposing BAs as a treatment for brain disorders are steadily increasing (Bhargava et al., 2020;Wu et al., 2020;Jin et al., 2021).

Interestingly, bariatric surgeries increase circulating BAs and, in mice, cause loss of preference for dietary fat (Scholtz et al., 2014;Bensalem et al., 2020) and cocaine (Reddy et al., 2018). Further, feeding BAs to mice recapitulated the effects of bariatric surgery on cocaine preference, strongly supporting their ability to modulate reward circuitries in the CNS as well as dopamine (DA) neurotransmission (Reddy et al., 2018). Understanding the molecular determinants of these actions could facilitate BA-based therapies for addiction as well as identify potential new drug targets.

BAs are directly derived from cholesterol and retain a high degree of structural similarity with the parent molecule. The importance of cholesterol in monoamine transporter (MAT) function and membrane localization is well documented (Foster et al., 2008;Chang and Rosenthal, 2012). Cholesterol depletion modulates both MAT expression (Foster et al., 2008;Gabriel et al., 2013) and activity (Magnani et al., 2004;Adkins et al., 2007;Foster et al., 2008;Hong and Amara, 2010). In addition, there are indications of direct interactions between cholesterol and MATs such as the DA transporter (DAT) and human serotonin transporter (hSERT) (Cremona et al., 2011;Jones et al., 2012;Penmatsa et al., 2013;Wang et al., 2015;Coleman et al., 2016;Zeppelin et al., 2018). Thus, investigations of the functional relevance of interactions between cholesterol-like molecules and MATs are needed to further define the central regulatory role of these sterols. This study demonstrates the ability of multiple BAs to alter DAT electrical activity and defines the structural changes underlying these alterations.

## 2 Materials and Methods

### 2.1 Solutions

Composition of buffered solution (ND96) (in mM): NaCl 96, KCl 2, CaCl_2_ 1.8, MgCl_2_ 1, HEPES 5, pH 7.6. Composition of NDE solution; ND96 plus 2.5 mM pyruvate and 50 μg/mL Gentamycin sulphate. Composition of external control buffer (ND98) (in mM): NaCl 98, MgCl_2_ 1, and CaCl_2_ 1.8 with or without 0.01% DMSO. In tetramethylammonium (TMA)-chloride “zero sodium” buffer (TMA98), equimolar TMA replaces NaCl. In Li^+^ buffer, equimolar lithium chloride replaces NaCl. The final pH was adjusted using respective hydroxides (NaOH or TMAOH or LiOH) to 7.6 for all external solutions. Substrates used were Dopamine (DA) (Calbiochem - Sigma, Milan, Italy), Lithocholic acid (LCA) (Sigma), and Obeticholic acid (OCA) (Adipogen, Switzerland). LCA or OCA powder was dissolved in DMSO at 50 mM and 100 mM, respectively, to generate stock solutions.

### 2.2 Oocytes collection and preparation

Oocytes were obtained from adult *Xenopus laevis* females. Animals were anaesthetised in 0.1% (w/v) MS222 (tricaine methanesulfonate; Sigma) solution in tap water. Abdomens were sterilized with antiseptic agent (Povidone-iodine 0.8%), laparotomy was performed, and portions of the ovary were collected. The oocytes were treated with 0.5mg/mL collagenase (Sigma Type IA) in ND96 calcium-free for at least 30 min at 18 °C. Healthy and fully-grown oocytes were selected and stored at 18 °C in NDE solution (Bossi et al., 2007). The day after the removal, the oocytes were injected with cRNA using a manual microinjection system (Drummond Scientific Company, Broomall, PA). Injected concentrations were 12.5 ng/50 nl for the mouse dopamine transporter (mDAT), 2 ng/50 nl for human Takeda G protein-coupled receptor (hTGR5).

### 2.3 cRNA preparation

mDAT cDNA in pcDNA3.1 was kindly gifted from Prof. Dr. Harald Sitte of Medical University of Vienna. The cDNA mDAT was amplified with forward and reverse primers containing SmaI and EcoRI restriction sites respectively (5’-GACT<u>CCCGGG</u>ACCCATGAGTAAAAGCAAATG-3’; 5’-GCAT<u>GAATTC</u>TTACAGCAACAGCCAATGGCGC-3’). The amplified coding sequence was then subcloned into the pGHJ vector after double digestion with SmaI e EcoRI restriction enzymes (Promega). hTGR5 gene was in pCMV6-Entry (GPBAR1 Human cDNA ORF Clone, NM_001077191; Origene Technologies, Inc., Rockville, MD, USA). The two plasmids were linearized with SalI (mDAT) and with NdeI (hTGR5), in vitro capped, and transcribed using T7 RNA polymerase. Enzymes were supplied by Promega Italia. The oocytes were incubated at 18°C for 2-3 days prior to electrophysiological experiments. The experimental protocol was approved locally by the Committee of the “Organismo Preposto al Benessere degli Animali” of the University of Insubria (OPBA-permit #02_15) and nationally by Ministero della Salute (permit nr. 1011/2015).

### 2.4 Electrophysiology

Electrophysiological studies were performed using the Two-Electrode Voltage Clamp (TEVC) technique (Oocyte Clamp OC-725B; Warner Instruments, Hamden, CT, USA). Controlling software was WinWCP version 4.4.6 (J. Dempster, University of Strathclyde, UK). Borosilicate microelectrodes, with a tip resistance of 0.5–4 MΩ, were filled with 3 M KCl. Bath electrodes were connected to the experimental oocyte chamber via agar bridges (3% agar in 3 M KCl). The holding potential was kept at −60 mV for all the experiments. The mean of the transport-associated currents plotted in the scatter diagrams were determined by subtracting the current recorded in the ND98 buffer from the current recorded in the presence of DA or OCA. In experiments using TMA, subtraction was performed with the current in TMA98 buffer. The current in the presence of OCA is the maximal current measured at the peak. To chelate intracellular calcium, oocytes were injected with 50 nl of an intracellular solution containing 13 mM EGTA 30 minutes prior to electrophysiological recording. Intracellular solution had the following composition (in mM): KCl 130, NaCl 4, MgCl_2_ 1.6, HEPES 10, glucose 5, pH 7.6.

### 2.5 Data analysis

Data analysis was performed using Clampfit 10.2 software (Molecular Devices, Sunnyvale, CA, USA, www.moleculardevices.com); OriginPro 8.0 (OriginLab Corp., Northampton, MA, USA, www.originlab.com) and GraphPad Prism (www.graphpad.com/scientific-software/prism) were used for statistical analysis and figure preparation.

### 2.6 Structural models of mDAT and hDAT

The mDAT (residues R56 to N596; UniProt ID O55192) in the outward-facing open (OF*o*), occluded, and inward-facing open (IF*o*) states were generated using SWISS-MODEL (Waterhouse et al., 2018) based on the structures resolved for OFo Drosophila DAT (dDAT) (PDB: 4M48) (Penmatsa et al., 2013), occluded hSERT (PDB: 6DZV) and IFo hSERT (PDB: 6DZZ) (Coleman et al., 2019). Homology models for hDAT in these conformational states were taken from previous work (Cheng et al., 2018;Aggarwal et al., 2021).

### 2.7 Docking simulations

OCA 3D structures were downloaded from the ZINC database (Irwin and Shoichet, 2005) (ZINC14164617) and DrugBank (Wishart et al., 2006) (DB05990). The net charge of OCA is indicated to be -1 in the ZINC database, and zero in DrugBank. Docking simulations were performed using both electronic states, designated as OCA(-) and OCA(n) for the negatively-charged and neutral OCAs, respectively. The binding sites and binding poses on both mDAT and hDAT, in different conformational states, were assessed using docking simulation software AutoDock4 (Morris et al., 2009) and AutoDock Vina (Trott and Olson, 2010). Autodock Vina simulations were carried out using a grid with dimensions set to 84 × 58 × 86 Å^3^ and center at (−1.22 Å, 1.16 Å, -6.26 Å) for each conformer and each transporter. This grid box encapsulated the entire structure of the transporter. The exhaustiveness of the simulation was set to 50 and the algorithm returned 20 binding modes of interest for each conformer. AutoDock4 simulations were performed following previously published protocols (Cheng et al., 2015;Aggarwal et al., 2021). Briefly, Lamarckian genetic algorithm with default parameters was employed with the maximal number of energy evaluations set to 2.5 × 10^7^. The binding energy was estimated from the weighted average of multiple binding poses at a given site observed in 100 independent runs.

## 3 Results

### 3.1 OCA induces a DAT-dependent, transient inward current

To begin to investigate the regulatory effects of OCA on DAT electrical activity, *Xenopus laevis* oocytes were tested by two-electrode voltage clamp (TEVC) (Fig. 1A). Perfusion of DA onto the oocytes expressing the mouse DAT (mDAT) generated an inward transport current (−12.5nA ± 0.63) (Fig 1B-C, left), confirming the expression and functionality of the transporter. On the same oocytes, following DA washout, perfusion of OCA also generated an inward transport current (−8.28nA ± 0.6), however, this membrane conductance exhibited a lower amplitude and inactivated rapidly (Fig 1B-C, left). The OCA-induced transient inward current was also elicited when OCA was perfused prior to DA exposure (data not shown). Oocytes co-expressing mDAT together with the human bile acid receptor, hTGR5, showed comparable current amplitudes in response to DA (−12.88nA ± 0.9) and OCA (−9.37nA ± 0.82) compared to oocytes expressing mDAT alone (Fig. 1B, center, and 1C, right). Further, OCA altered the membrane conductance only in oocytes expressing mDAT; no currents were observed from expression of hTGR5 alone (Fig. 1B, right and 1D). This data suggests the inward current induced by OCA is directly mediated by mDAT.

**Figure 1:**
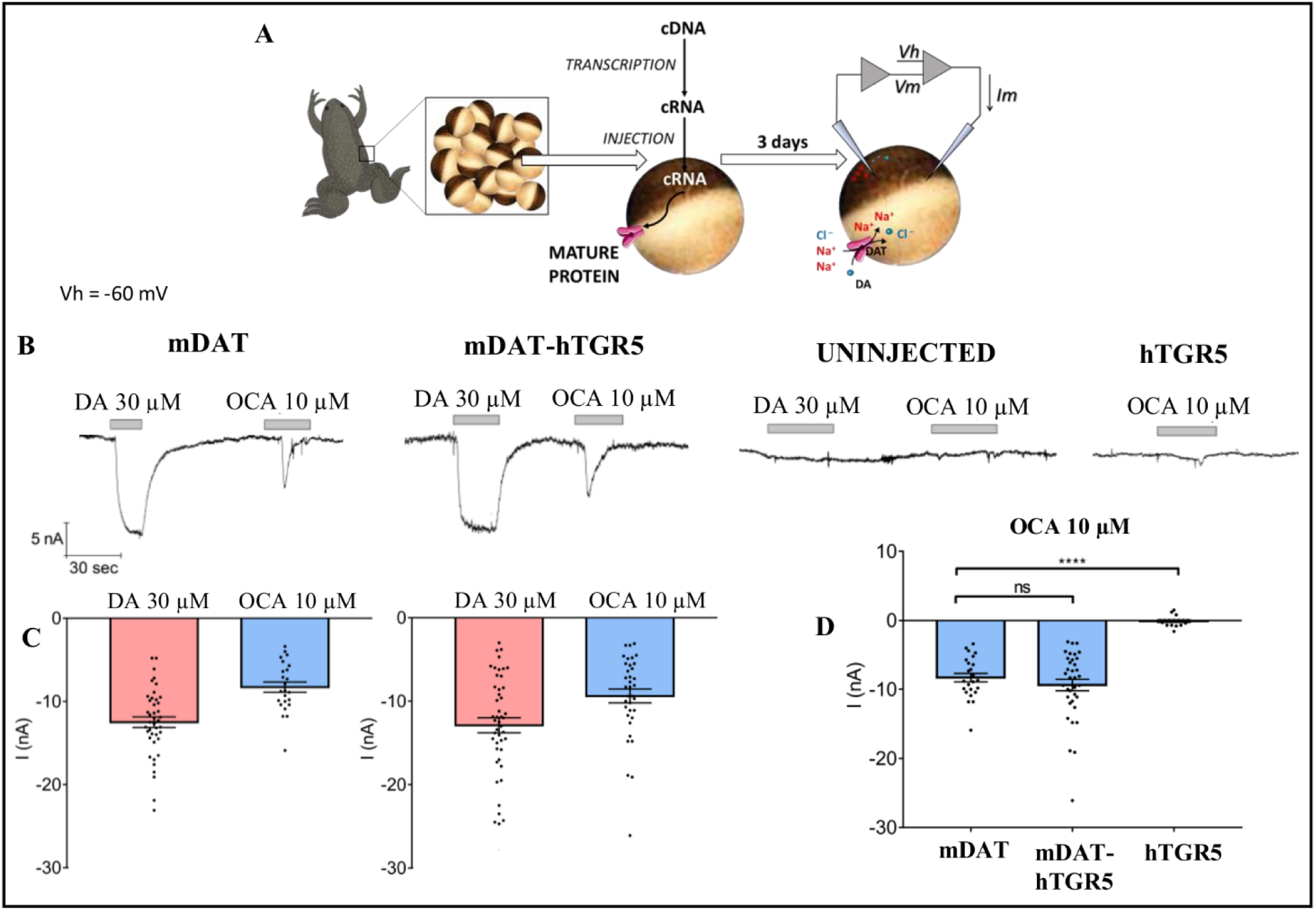
OCA generates an electrical current through the dopamine transporter. **(A)** Schematic representation of oocyte collection, cRNA synthesis and injection, and TEVC technique. **(B)** Representative traces of currents recorded by TEVC (Vh= -60 mV) from oocytes expressing mDAT, without or with hTGR5, uninjected, or expressing hTGR5 alone. The oocytes were perfused with 30 μM DA or 10 μM OCA. **(C)** Mean of the maximal DA-associated and OCA-induced currents (I nA ± SE of 14–47 oocytes, 6 –11 batches) in oocytes expressing mDAT (C, left) or mDAT plus TGR5 (C, right). **(D)** Mean of the maximal OCA-induced currents in oocytes expressing the proteins indicated. **** p<0.0001; one-way ANOVA followed by Tukey’s multiple comparison test (DF= 2 between columns).

Several members of the SLC6 family exhibit basal leak currents (Lester et al., 1994;Bossi et al., 1999;Andrini et al., 2008) that are augmented in the presence of Li^+^ ions. The Li^+^ leak current can be utilized to highlight the effect of molecules that interact with transporters and modify their electrical activity in the absence of Na^+^. DAT shows a significant Li^+^ leak current that is partially blocked upon DA perfusion (Giros et al., 1992;Sonders et al., 1997). Therefore, we used Li^+^ to investigate whether OCA, similarly to DA, binds mDAT in the absence of Na^+^. Oocytes were perfused first either with DA or OCA in Na^+^ bathing buffer (Fig. 2A-B). Switching to Li^+^ bathing buffer induced a large inward current (−92.41nA ± 7.45) (Fig. 2A-B). As expected, the addition of DA to the Li^+^ bathing buffer partially blocked the Li^+^-leak current (Fig. 2A-B). This inhibition occurred in two phases; a rapid transient component (current at the peak: -25.72nA ± 4.07) followed by a steady-state condition (−38.97nA ± 5.11). After DA removal, the Li^+^ leak current returned to initial values (Fig. 2A). Interestingly, perfusion of OCA also partially blocked the Li^+^ leak current (Fig. 2A-B). As with DA, this inhibition displayed two phases; a transient rapid component (current at the peak: -66.86nA ± 6.13) and a steady-state component (−82.42nA ± 7.69). Inhibition of the Li^+^ leak current is a strong indication of direct binding of OCA to the transporter in the absence of Na^+^.

**Figure 2:**
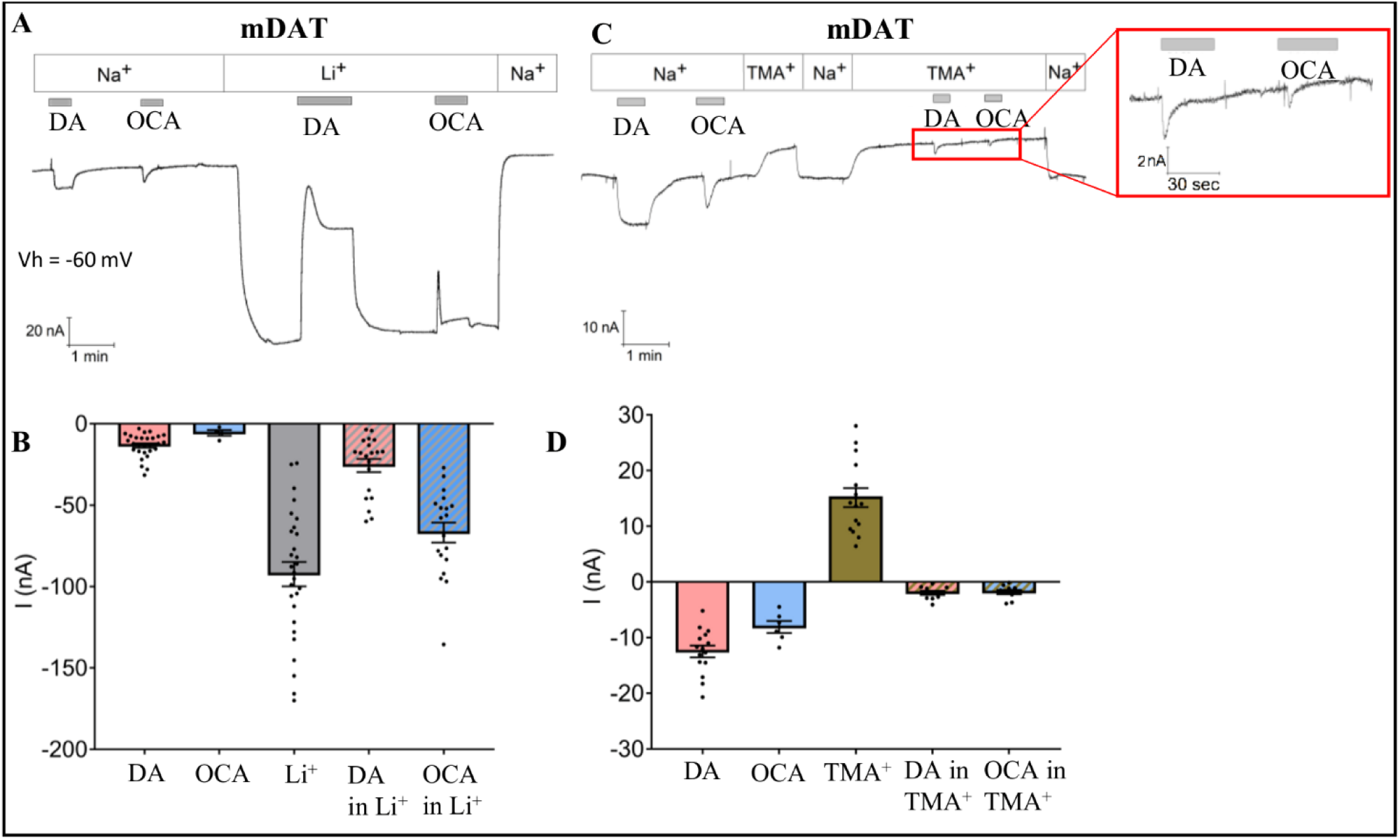
OCA modulates the mDAT-mediated leak current. **(A)** Representative current traces recorded with TEVC (Vh= -60 mV) in oocytes expressing mDAT perfused with 30 μM DA or 10 μM OCA in ND98 or Li98 buffer. **(B)** Mean of maximal currents under conditions shown in A (I nA± SE of 6-28 oocytes, 4 batches). **(C)** Representative trace from oocytes expressing mDAT perfused with 30 μM DA or 10 μM OCA in ND98 or TMA98 buffer. **(D)** Mean of maximal currents under conditions shown in C (I nA± SE of 6-15 oocytes, 3 batches).

### 3.2 mDAT-mediated OCA current is Na^+^ dependent

In addition to a coupled mechanism, Na^+^ also permeates through DAT in the absence of DA. This generates a leak current that can be uncovered when Na^+^ is substituted by non-permeant cations such as Choline or TMA^+^ (Sonders et al., 1997). To better understand the effect of OCA on membrane conductance and the relevance of Na^+^ in the OCA-induced transient inward current, experiments were repeated with TMA^+^ as the cation substituting for Na^+^ in the bathing buffer. TMA^+^ blocked the Na^+^-leak current (15.12nA ± 1.71) (Fig. 2C-D). As expected, in the presence of TMA^+^, DA elicited only a fast-transient inward current (−1.98nA ± 0.37), confirming that Na^+^ is necessary for mDAT-mediated DA currents. OCA behaved similarly to DA, the transient inward current was still present, but significantly reduced in amplitude (−1.83nA ± 0.38) (Fig. 2C-D).

### 3.3 OCA current is not due to an increase of intracellular calcium

OCA has been shown to induce intracellular Ca^2+^ fluctuations (Hao et al., 2017). Thus, it is possible that the OCA-induced transient inward current could be generated by the activation of chloride conductance due to an increase in intracellular Ca^2+^ concentrations. To investigate this possibility, experiments were conducted in oocytes expressing mDAT and injected with the Ca^2+^-chelating agent EGTA (Fig. 3A). The presence of EGTA did not alter mDAT-mediated currents elicited by DA or OCA (Compare Fig. 3B with Fig. 1B-C). Maximal OCA-induced transient inward currents were also not altered by the presence or absence of EGTA (Fig. 3C). Together, these data strongly suggest that intracellular Ca^2+^ does not regulate OCA-induced currents and further suggest that effects of OCA are mediated by direct interaction with mDAT.

**Figure 3:**
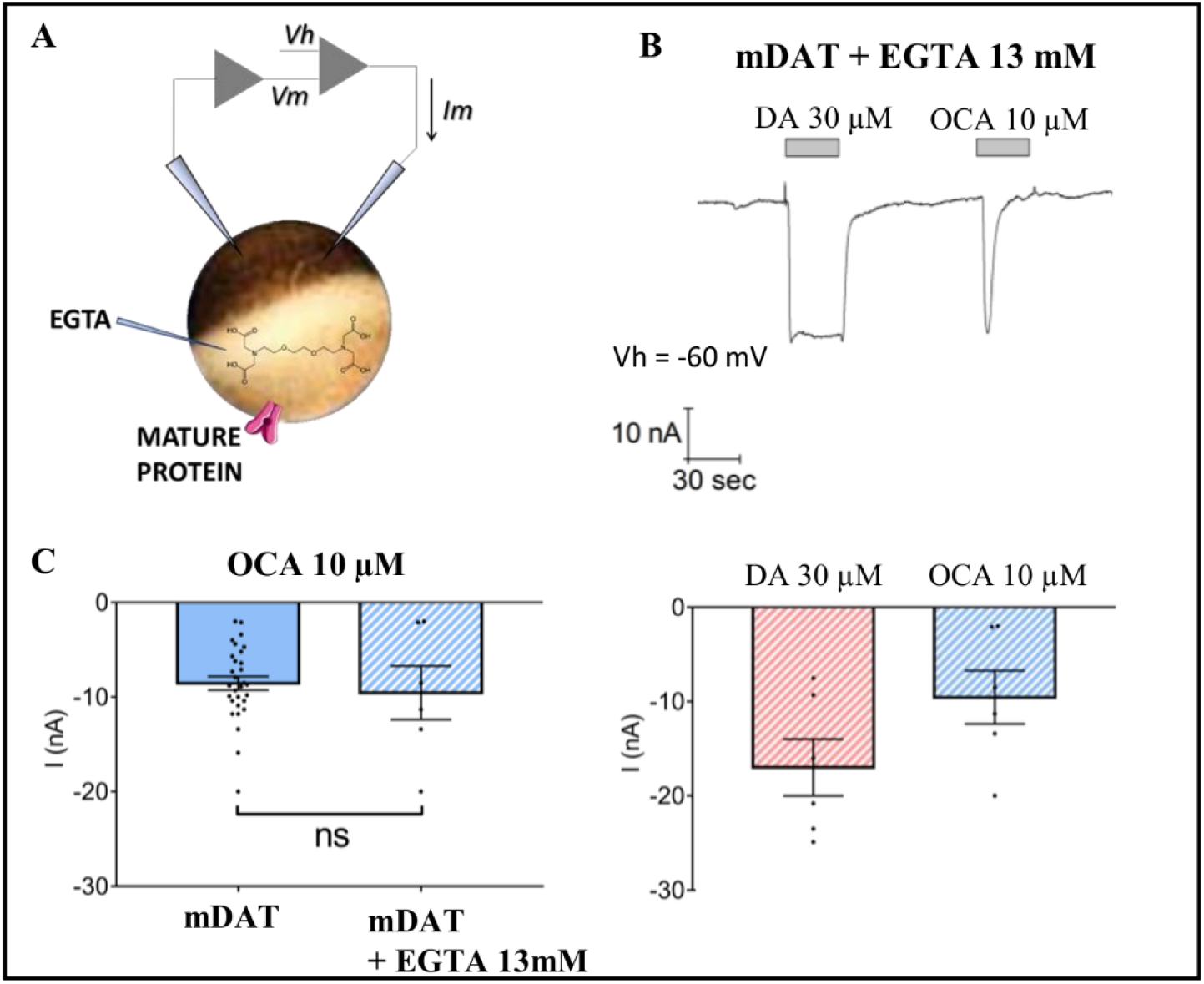
Intracellular calcium does not regulate OCA-induced currents. **(A)** Schematic representation of the EGTA injection technique. **(B)** Representative trace of current recorded by TEVC (Vh= -60mV) in oocytes expressing mDAT and injected with 13 mM EGTA in intracellular solution 30 minutes before exposure to 30 μM DA or 10 μM OCA in ND98 buffer (top) and the mean of maximal transport-associated and OCA-induced transient currents (bottom) (I nA± SE of 6-7 oocytes, 2 batches). **(C)** Mean of maximal OCA-induced transient currents in mDAT oocytes with or without injection of EGTA (n=7 and 24; p>0.05 by Student’s t-test).

### 3.4 OCA does not alter either DAT-mediated DA currents or DA affinity

Data thus far indicate that OCA interacts directly with mDAT in the absence of DA. We next investigated whether OCA regulates DAT-mediated DA-induced currents. Currents generated from increasing concentrations of DA where unaltered by the presence of OCA. (Fig. 4A). Maximal currents were fitted to a Hill equation (Fig. 4B). These data indicated that the presence of OCA does not significantly affect either the affinity or the maximal transport currents.

**Figure 4:**
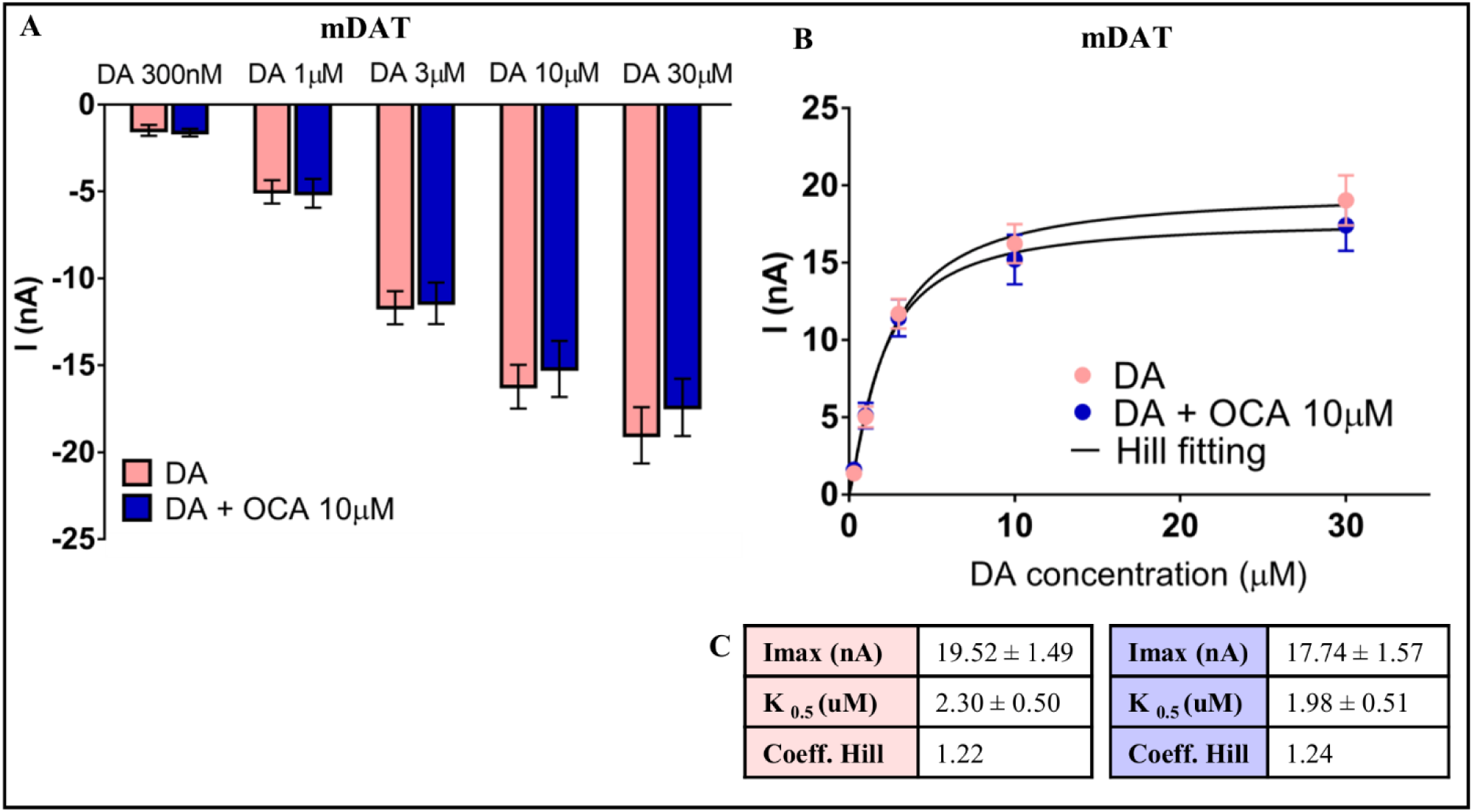
OCA effect on dopamine transport associated current. **(A)** Mean currents recorded at increasing concentrations of DA, as indicated, in the absence or presence of 10 μM OCA (I nA ± SE of 15-18 oocytes, 3 batches) (n=18; p>0.05 by Student’s t-test). **(B)** Data from A were fitted to a Hill equation. **(C)** Imax, K_0.5_, and Hill coefficient obtained from the fitting to Hill equation of the data represented in B (absence of OCA (*pink, left*); presence of OCA (*blue, right*)).

### 3.5 Lithocholic acid induces DAT-mediated currents

To determine whether the transient current generated through mDAT is specific to the synthetic bile acid OCA, or is a common phenotype induced by bile acids, we investigated the effect of the natural bile acid, lithocholic acid (LCA). OCA and LCA share the same sterol-based structure with differing R groups at positions 5 and 6 of the B ring (Fig. 5A). Specifically, ethyl and hydroxyl groups present in OCA are substituted by hydrogen in LCA. Similar to OCA, perfusion of LCA onto mDAT-expressing oocytes induced a transient inward current (−10.25nA ± 2.06) (Fig. 5B-C). Direct comparison of OCA and LCA revealed no significant differences in their ability to induce DAT-mediated currents (Fig. 5C). These data suggest that the ability of OCA to promote DAT-mediated currents is shared by another BA.

**Figure 5:**
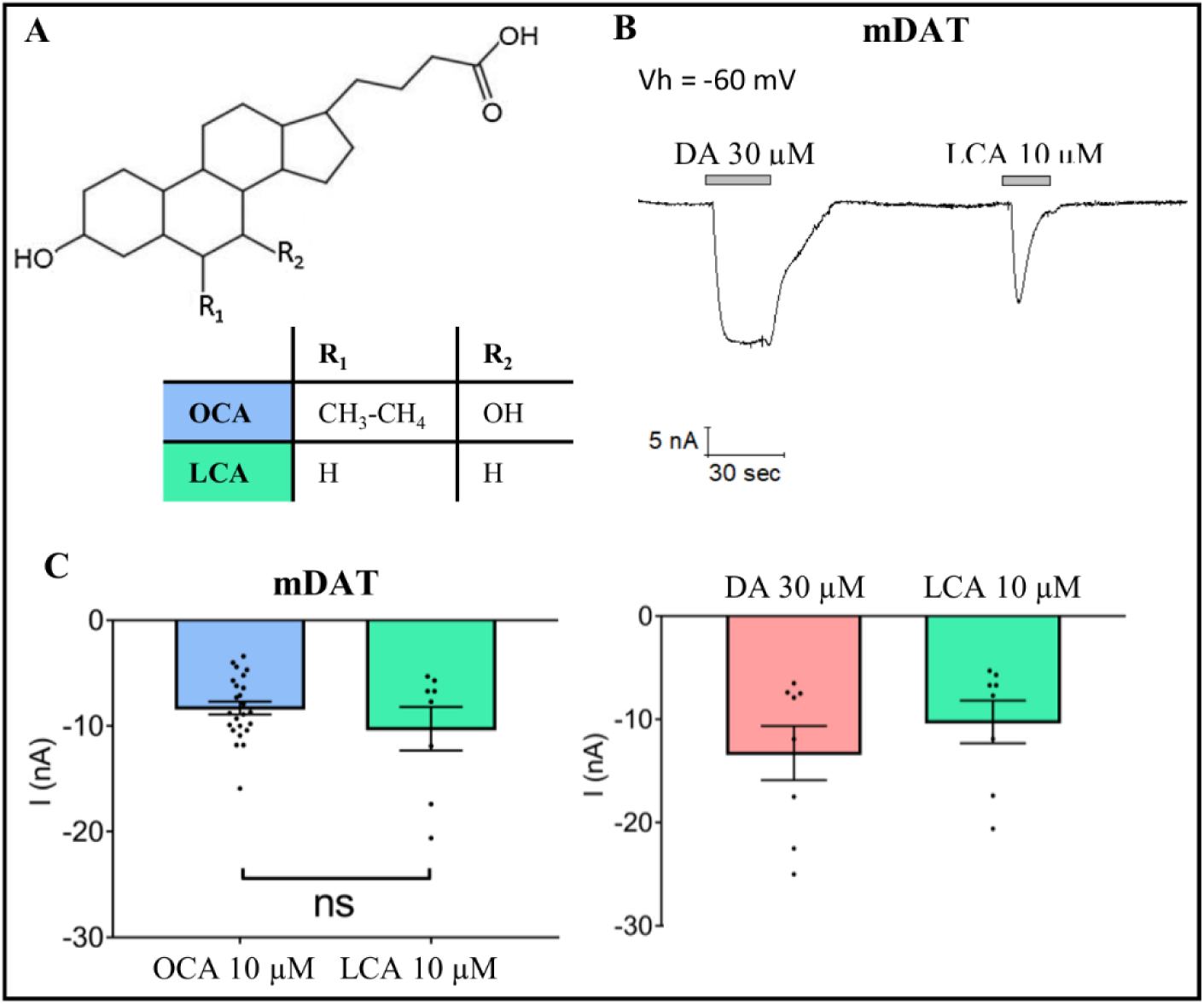
LCA effect on Dopamine Transporter. **(A)** Structure of OCA and LCA **(B)** Representative trace of current recorded with TEVC (Vh= -60mV) in oocytes expressing mDAT and perfused with 30 μM DA or 10 μM LCA in ND98 buffer (top). Mean of maximal dopamine transport-associated and LCA-induced transient currents (I nA ± SE of 8 oocytes, 3 batches) (bottom). **(C)** Mean of maximal transient currents elicited by OCA or LCA in mDAT-expressing oocytes (n=8; p>0.05 by Student’s t-test).

### 3.6 The binding pose of OCA(n) suggests that it binds to DAT in a non-competitive, drug accommodating fashion, that allows simultaneous binding of DA

These data demonstrate that at least two BAs can bind to mDAT. To identify potential sites for binding of OCA onto mDAT and hDAT, docking simulations were performed, using AutoDock4 (Morris et al., 2009) and AutoDock Vina (Trott and Olson, 2010). To account for alternative protonated and deprotonated states of OCA, both the neutral (OCA(*n*)) and negatively-charged (OCA(-)) forms (Fig. 6D) were used in the simulations. Computations were performed for the outward-facing *open* (OF*o*), *occluded* (OCC), and inward-facing *open* (IF*o*) states of both mDAT and hDAT. The computations revealed that OCA selected similar binding poses, with comparable binding affinities, for either transporter when the same conformational state was targeted regardless of the species. Therefore, representative results for hDAT are presented unless otherwise stated.

**Figure 6.**
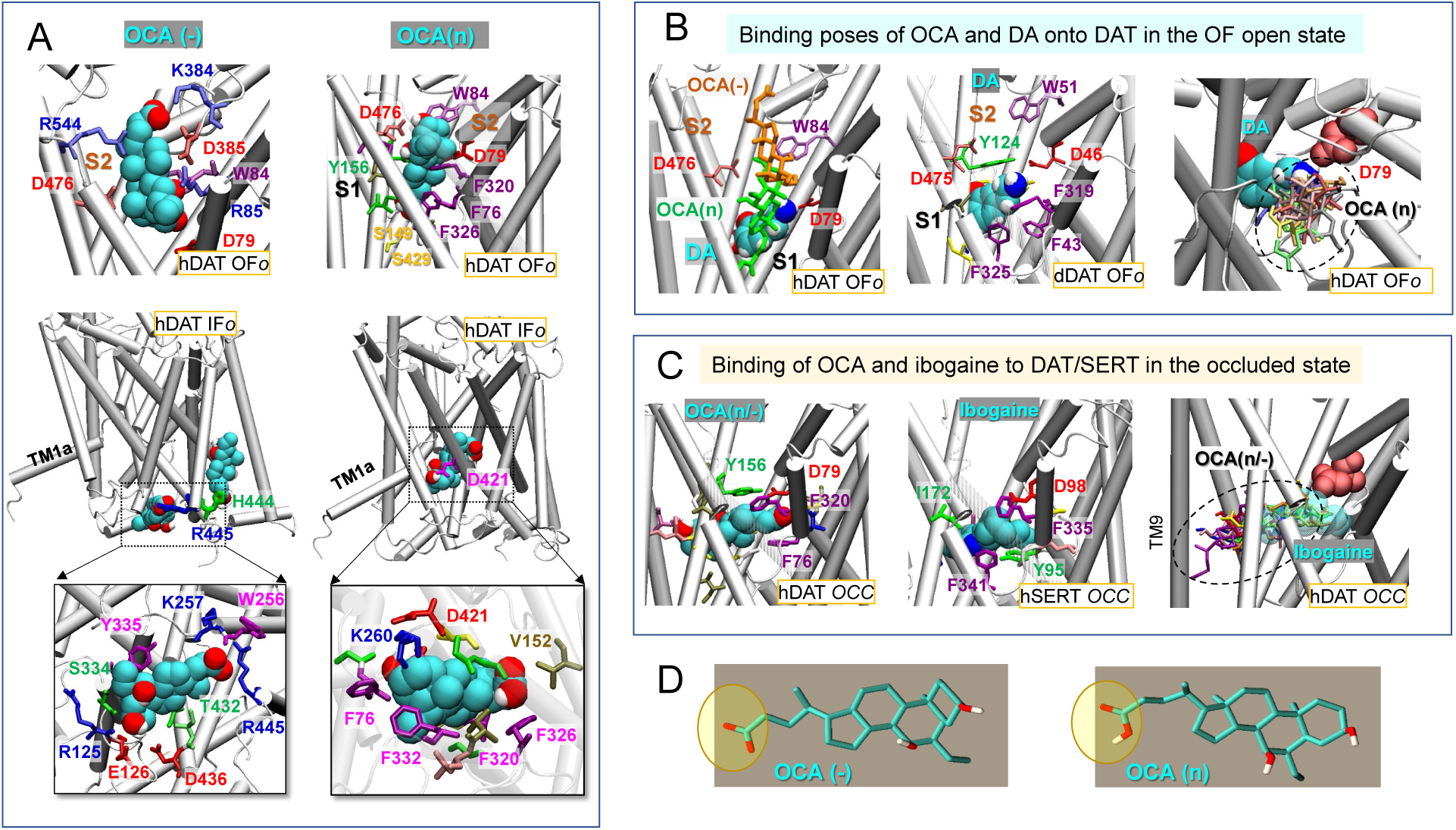
OCA binding to DAT depends on its protonation state and DAT conformation. **(A)** Binding of negatively charged (*left*) and neutral (*right*) OCA (cyan) to hDAT in the OFo (top two panels) and IFo (*middle and bottom panels*) states. The binding energies to OFo hDAT are -6.1 kcal/mol (OCA(-)) and -9.9 kcal/mol (OCA(n)); and those to the IFo hDAT are -7.5 kcal/mol (OCA(-)) and -8.5 kcal/mol (OCA(n)). Interacting residues making atom-atom contacts closer than 4 Å with OCA are shown in panels A-C. **(B)** Comparison with the known DA-binding pose. Alignment of the top binding poses of OCA(n) (*green sticks*), OCA(-) (*orange sticks*) (*left panel*) and the resolved DA (van der Waals (vdW) format) bound dDAT (PDB: 4XP1) (*middle panel*), and detailed view of DA-binding site and comparison of the non-overlapping spaces occupied by OCA (multiple binding poses) and DA (*right panel*). **(C)** Top 1 binding pose of OCA(n/-) (vdW format; -10.2 kcal/mol) to hDAT occluded (OCC) conformer (*left panel*), the resolved ibogaine-bound to hSERT in the OCC state (PDB:6DZV) (*middle panel*) and alignment, onto hDAT, of top 10 binding poses of OCA(n) (*sticks in different colors*; average = -9.1±1.0 kcal/mol) and ibogaine bound to SERT (transparent vDW) (*right panel*). OCA, DA and ibogaine are shown in van der Waals (vdW) format in A-C, with cyan, red, blue and white spheres representing carbon, oxygen, nitrogen and hydrogen atoms. **(D)** Molecular structures of OCA in two protonation states.

Figure 6A illustrates the top-ranking distinct binding poses and sites of both OCA forms, observed in docking simulations, onto hDAT in OF*o* and IF*o* states, panel B shows the comparison to the DA-bound structures resolved for hDAT and dDAT (Wang et al., 2015), and panel C to ibogaine-bound hDAT and hSERT (Coleman et al., 2019). Notably, in the case of the OF*o* conformer, the most favorable binding sites for both the neutral and negatively charged forms of OCA are within the extracellular (EC) vestibule but the exact locations are determined by the protonation state of the ligand. OCA(-) mainly occupies the S2 site proposed to allosterically modulate transport (Cheng and Bahar, 2019), whereas OCA(n) binds an extended region spanning between the primary (S1) and secondary (S2) sites (Fig. 6A, top panels). The binding pose of OCA(-) thus differs from that of the DA bound to OF*o* DAT (Fig. 6B) where the residue equivalent to hDAT D79 (D46 in dDAT) plays a major role in coordinating the binding of the amine group through salt bridge formation (Penmatsa et al., 2013;Wang et al., 2015;Cheng and Bahar, 2019). Notably, the top binding poses of OCA do not block the binding of substrate DA (Fig. 6B, left panel), such that DA is able to bind in the proximity of OCA. This is consistent with the non-competitive binding of OCA revealed in TEVC experiments (Fig. 4).

For the IF*o* conformer, no high affinity binding is observed to the EC vestibule. For OCA(n), the top binding site is in proximity to D421 within the intracellular (IC) vestibule (Fig. 6A, bottom right panel). OCA(-) preferentially binds near the IC entrance, in the proximity to R445, and no high affinity binding is observed within the vestibule. Notably, docking simulations also revealed binding instances within the transmembrane region, including the site known to be occupied by cholesterol near TM1a (Penmatsa et al., 2013;Cheng and Bahar, 2019), or a site close to residues R443/H444 (Fig. 6A, middle left panel) reported previously as a PIP_2_ binding site (Belovich et al., 2019). However, these sites could potentially be obstructed by lipid molecules, which are not included in docking simulations, and hence will not be further elaborated.

Unexpectedly, in the case of *occluded* DAT conformer (Fig. 6C) both charged and neutral OCAs (OCA(*n*/-)) are predicted to bind to the S1 site, with almost identical affinity, and the binding pose closely resembles that resolved for ibogaine-bound hSERT. Of note, the binding pocket in the occluded state may be much larger than anticipated, with possible involvement of TM9 (Fig. 6C, right panel).

Taken together, these docking simulations suggest that the binding sites and affinities depend on the net charge carried by OCA and on the conformational state of the transporter. In the OF*o* state, the binding site for OCA(*n*) closely neighbors, but does not overlap, the S1 site resolved for DA (Wang et al., 2015); whereas, the binding site of OCA(-) overlaps with the broadly-defined S2 site (Fig 6A-B). Notably, in the occluded state of the transporter, OCA appears to select a binding site similar to that resolved for ibogaine-bound to hSERT (Coleman et al., 2019) regardless of its charge. In the IF*o* state, OCA binds near D421 or R445, depending on its protonation state.

## 4 Discussion

These current findings highlight previously undocumented interactions and functional impact of BAs on DAT function. Specifically, we show that this interaction promotes a transient current that is Na^+^ dependent. Notably, OCA is capable of inhibiting, as observed for DA (Sonders et al., 1997), a DAT-mediated Li^+^-leak current suggesting that BAs can interact with DAT even in the absence of Na^+^. The transient nature of the DAT-mediated OCA currents suggests that OCA, although capable of initially gating charge movement, ultimately induces an occluded conformation of the transporter. Furthermore, the OCA-induced current does not depend upon changes in intracellular calcium fluctuations nor TGR5 signaling.

It is important to consider the possibility that the transient current gated by OCA is a DAT-mediated Na^+^-dependent leak current. Indeed, Na^+^ substitution with TMA^+^ strongly reduces the ability of OCA to induce this current. In previous *in silico* study of DA-free DAT conformers (Cheng et al., 2018;Cheng and Bahar, 2019), the EC- and IC-exposed helices were not as tightly packed as in the *occluded* DA-bound form. These conformers occasionally gave rise to simultaneous opening of both the EC and IC gates such that an intermittent formation of a water channel was detected. Notably, the sodium permeation path (Aguilar et al., 2021) coincides with that of water channeling (Cheng et al., 2018;Cheng and Bahar, 2019). This path, observed *in silico*, may also be associated with DAT-mediated ion fluxes or leak currents (Ingram et al., 2002;Erreger et al., 2008). Together with previous simulations (Cheng and Bahar, 2015), the current ionic substitution experiments show that DA-binding and translocation induce a tightening in intramolecular interactions to block current leakages.

DA successively binds S2 and S1 sites, stimulates extracellular (EC) gate closure upon stabilization in the site S1, and cooperatively restores the compact association of the EC-vestibule while translocating to the intracellular (IC) vestibule (Cheng and Bahar, 2015). The IC opens to the cytoplasm only after compaction/closure of the EC-exposed region (Cheng and Bahar, 2015). Likewise, OCA binding to the S2 substrate site in the charged state and deeper insertion and translocation after protonation may induce similar intramolecular rearrangements to block the leak currents, as observed in the presence of Li^+^. Differences were observed between the effects of OCA and DA on DAT conductance, in terms of steady-state currents. This may be due to the fact that release of DA and simultaneously-bound Na^+^ restores DAT into a transporter mode (Borre et al., 2014), whereas the non-substrate OCA, will not facilitate full procession to this state resulting in a current that is transient.

Li^+^ leakage in DAT is dependent on the Na2 site rather than the Na1 site (Borre et al., 2014). D421 in hDAT coordinates the binding of the Na^+^ ion to the Na2 site (Borre et al., 2014;Cheng and Bahar, 2015). Notably, the current docking simulations also indicate that D421 may contribute the binding of OCA(n) (Fig. 6A) in the IF*o* state. The direct binding of OCA to D421 may thus potentially block Li^+^ permeation. However, it is not possible to rule out that the inhibition of Li^+^ current by OCA may reflect the shift of conformational equilibrium between different functional states along the transport cycle, as proposed for DA (Borre et al., 2014).

The docking simulations also provide important insights on the potential functional relevance of the OCA-DAT interaction. Surprisingly, these results suggest that OCA binds a similar site to that resolved for ibogaine-bound hSERT (Coleman et al., 2019) in the *occluded* state (see Fig. 6C). Ibogaine is a non-competitive inhibitor for both DAT and SERT and has been proposed to stabilize the transporters in the IF conformation (Jacobs et al., 2007;Bulling et al., 2012). Given that OCA is observed to bind in a non-competitive way with respect to DA (Fig. 4), the binding of OCA to DAT, in the absence of DA, may stabilize DAT in the occluded or IF*o* state, thus interfering with the transport cycle.

A separate site for OCA on DAT, of equal potential functional impact, is facilitated through hDAT R445 and D436 (Fig. 6A). This binding may stabilize the IF*o* state by preventing the re-formation of the IF gating salt bridges, R60-D436 and R445-E428, that are required for DA transport (Cheng and Bahar, 2019). Recently, molecular modeling found that the infantile Parkinsonism-Dystonia associated substitution, R445C in hDAT, disrupted a phylogenetically conserved intracellular network of interactions and promoted a channel-like intermediate of hDAT. These rearrangements lead to the permeation of Na^+^ from both the EC and IC solutions (Aguilar et al., 2021).

Lastly, the findings reported in Figure 5 confirm that binding to DAT is not unique to OCA, but takes place also upon application of LCA, a natural bile acid. This prompts the hypothesis that this class of molecules has an unforeseen regulatory potential on transporter function. BAs may represent novel pharmacological tools or candidate therapeutics for the treatment of diseases associated with DAT dysfunction.

## 7 Author Contributions

T.R., D.Z., M.H.C., and B.S. performed experiments, analysed data, prepared figures and contributed to writing the manuscript. A.M.C. contributed data analysis, writing and editing the manuscript. A.G., I.B., and E.B. designed and supervised the studies.

